# PEG-free, Triphosphate-Stabilized LNPs Enable Potent RNA Delivery

**DOI:** 10.64898/2025.12.19.695515

**Authors:** Gholan Hamraz, Mario A. Saucedo-Espinosa, Kerstin C. Walzer, Florian Frohns, Elfi Katrin Schlohsarczyk, Jakob Käppler, Björn C. Fröhlich, Stefan Schmidt, Carsten Hopf, Steffen Panzner

## Abstract

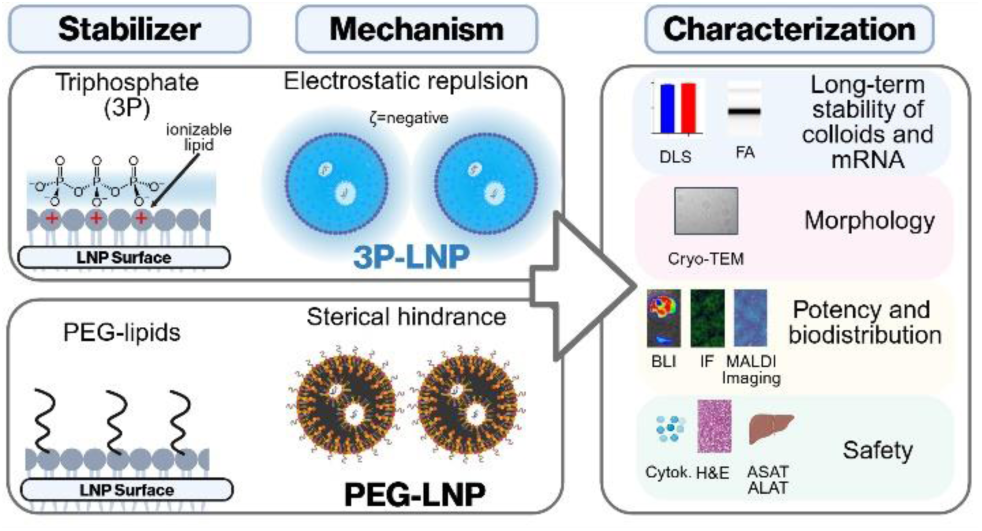

Polyethylene glycol (PEG)-lipids have long enabled lipid nanoparticle (LNP) formulations by providing steric stabilization and prolonged circulation. However, anti-PEG immune responses may impede repeated systemic dosing in mRNA therapeutics due to the phenomenon known as accelerated blood clearance (ABC). Here, we introduce a PEG-free, polyanionic alternative in which sodium triphosphate (3P) electrostatically associates with ionizable surface lipids, conferring long-term colloidal stability through charge repulsion rather than steric shielding. 3P-LNPs maintained size uniformity, morphology, and mRNA integrity for over nine months at 4 °C. In a proof-of-concept study involving a single intravenous injection of LNPs in mice carrying a luciferase and a GFP transgene, early expression was approximately two-fold higher compared to PEG-LNPs, while expression levels at 24 hours and hepatic tolerability remained comparable. Serum enzyme and cytokine profiles indicated no necrosis or inflammation. These findings establish 3P-LNPs as a stable, biocompatible platform free of PEG, with potential for repeat systemic mRNA administration while avoiding PEG-related effects.

---

The clinical success of mRNA therapeutics has been enabled by utilizing lipid nanoparticles (LNPs) delivery systems [1]. Clinically approved (LNPs) have four core components: an ionizable lipid, which facilitates mRNA encapsulation and release; two structural lipids, distearoylphosphatidylcholine (DSPC) and cholesterol; and a polyethylene glycol (PEG)-conjugated lipid [2] —commonly referred to as a stealth lipid—that modulates particle size [3], enhances colloidal stability by preventing aggregation, and extends circulation half-life [4]. PEG-lipids are not biologically inert when incorporated into LNPs, as they can potentially trigger anti-PEG antibody responses [5], which are associated with the accelerated blood clearance (ABC) [6] phenomenon, particularly after repeated systemic administration thereby reducing LNP delivery to target cells. Several PEG alternatives are being evaluated [7], including brushed-shaped polymers [8], cleavable PEG [9], and hydrophilic substitutes such as polysarcosine [10] or poly(vinyl pyrrolidone) [11]. While there is some evidence of reduced ABC risk and improved tolerability in preclinical models [12] [13], no alternative stealth-lipid-free LNP formulation has yet demonstrated full functionality in systemic *in vivo* mRNA-delivery. We hypothesized that multivalent anions could serve as an alternative to PEG. When bound to ionizable lipids on the LNP surface, anions such as sodium triphosphate (3P) is hypothesized to render the LNP stable through electrostatic repulsion. We comprehensively evaluated the physicochemical characteristics of 3P-stabilized LNPs (3P-LNP), including colloidal stability and mRNA encapsulation efficiency and integrity. Additionally, we assessed their *in vivo* performance and biodistribution in mice using a luciferase (Luc) and green fluorescent protein (GFP) transgene, liver enzyme profiles, cytokine responses, immune cell uptake, and histological outcomes, and investigated ionizable lipid distribution in the liver using matrix-assisted laser desorption/ionization mass spectrometry imaging (MALDI-MS Imaging). This systematic evaluation provides insights into stability and efficacy and safety of PEG-free formulations as a novel LNP platform technology.

To first investigate the cause of colloidal instability in PEG-free LNPs, we analyzed their ζ-potential across different pH conditions. Unstabilized LNPs (lacking both PEG and 3P) can successfully be produced and processed; however, they formed an instable colloid that aggregated within 48 hours when dialyzed against a physiological buffer. This coincides with a near-neutral ζ-potential at pH 7.4 (Fig. 1a), suggesting that insufficient electrostatic repulsion may underlie the lack of colloidal stability. PEG-free LNPs displayed a positive ζ-potential at pH 5.0, consistent with protonation of ionizable lipids, and a negative ζ-potential at pH 8.5 when the lipids are largely unprotonated, and some surface-exposed RNA might generate the charge instead. At pH 7.4, both the lipid and RNA charges cancel out each other, resulting in a lack of electrostatic repulsion and, eventually, aggregation. Supporting this view, ^1^H surface-proton NMR spectra at pH 7.4 revealed lipid signals accessible to the aqueous phase (Supplementary Figure S1), indicating that ionizable lipids are exposed at the particle interface. We hypothesized that multivalent anions having a high charge density, such as 3P, can bind to the surface-exposed cationic lipids, forming a negatively charged LNP starting at 5 mM (Fig. 1b). This additional anionic layer would provide an alternative means to electrostatic repulsion. Consistent with this mechanism, PEG-free LNPs supplemented with 5 mM 3P remained transparent at pH 7.4 after 48 hours at 4 °C, comparable to PEG-containing controls, whereas unstabilized LNPs (lacking both PEG and 3P) aggregated visibly under identical conditions (Fig. 1c).

**Figure 1:**
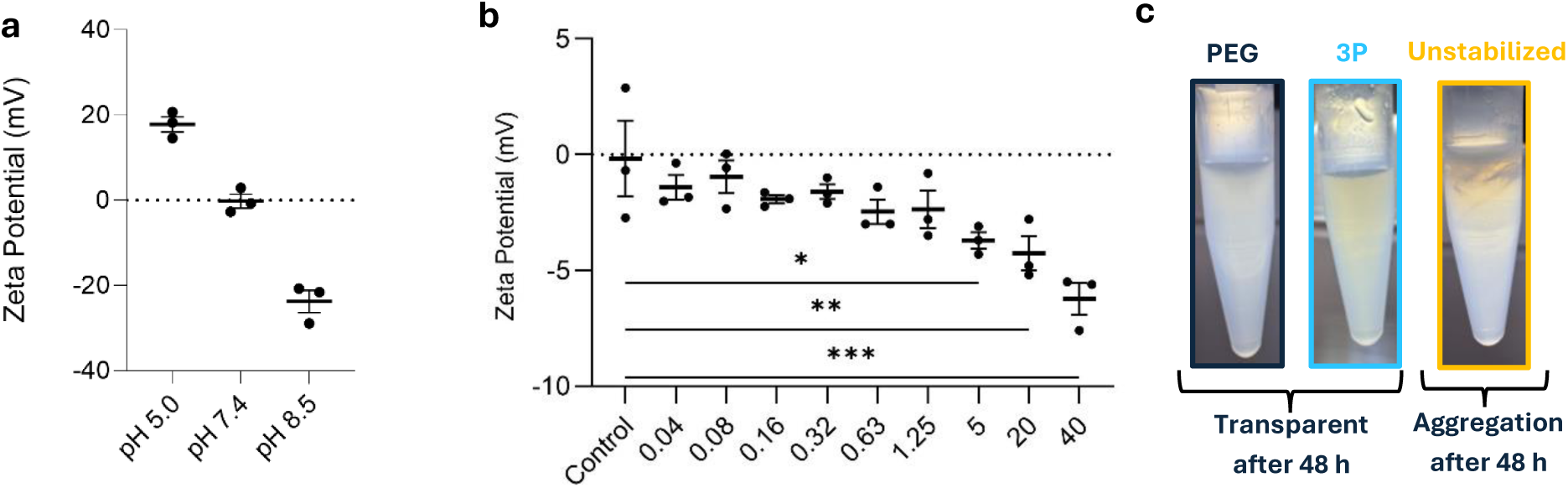
ζ-potential modulation and colloidal stability of unstabilized (lacking both PEG and 3P) and stabilized (with PEG or 3P) formulations. (a) ζ-potential measurements of unstabilized LNPs at varying pH levels (5.0, 7.4, and 8.5) in their native buffers. The ζ-potential transitions from positive at pH 5.0 (due to protonated ionizable lipids) to neutral at pH 7.4 (balance between ionizable lipids and negatively charged RNA), and negative at pH 8.5 (deprotonated ionizable lipids with RNA dominating surface charge). Data are presented as mean ± s.e.m. from three independent experimental replicates (n=3). (b) Modulation of ζ-potential of PEG-free LNPs by increasing concentrations of 3P (0–40 mM) in 10 mM HEPES buffer (pH 7.4). Higher 3P concentrations reduced the ζ-potential toward –7 mV, neutralizing residual protonated surface patches. Data are presented as mean ± s.e.m. from three independent experimental replicates (n=3). Statistical significance was determined using one-way ANOVA followed by Dunnett’s multiple comparisons test to compare all experimental groups to the control group. (c) Colloidal stability of PEG-LNPs, 3P-LNPs, and unstabilized LNPs at pH 7.4 after 48 h at 4 °C. PEG-LNPs and 3P-LNPs remained transparent, indicating preserved colloidal stability, while unstabilized LNPs showed visible aggregation. Statistical significance was defined as follows: (p > 0.05), * indicates significance (p ≤ 0.05), and ** indicates high significance (p ≤ 0.01). For clarity, statistically non-significant comparisons were not shown. Abbreviations: 3P, triphosphate; HEPES, 4-(2-hydroxyethyl)-1-piperazineethanesulfonic acid; h, hour; LNP, lipid nanoparticle; PEG, polyethylene glycol; S.e.m, standard error of the mean; ζ-potential, zeta potential.

To evaluate whether 3P can colloidally stabilize LNPs, we prepared three distinct formulations: conventional PEG-LNPs, 3P-LNPs without PEG, but stabilized by addition of 5 mM 3P during dialysis, and unstabilized LNP (Fig. 2a). All particles were produced with the ionizable lipid di(heptadecan-9-yl) 3,3’-((2-(4-methylpiperazin-1-yl)ethyl)azanediyl)dipropionate (BHD-C2C2-PipZ) (Supplemental Fig. S2), cholesterol, DSPC and optionally a PEG-lipid via a standard ethanol injection protocol (Fig. 2b). Uniform ∼80 nm particles with a polydispersity index (PDI) ≤ 0.1 and encapsulation efficiencies above 99% were produced (Fig. 2c, left and middle). Absence of PEG resulted in a ζ-potential of ∼0 mV for PEG-free LNP while 3P-LNP resulted in an intermediate ζ-potential of ∼−4 mV (Fig. 2c, right), indicating the binding of multivalent anions. For comparison, PEG-LNP exhibited a zeta potential of –8 mV, highlighting the chemical differences between PEG-LNP and PEG-free LNP formulations. Cryogenic transmission electron microscopy (Cryo-TEM) confirmed intact, spherical morphologies for PEG- and 3P-LNPs (Fig. 2d). Unstabilized LNPs were excluded from analysis due to aggregation. LNPs containing 5 mM monophosphate or citrate were additionally formulated; however, these formulations failed to maintain colloidal stability beyond five days. (Supplemental Fig. S3).

**Figure 2:**
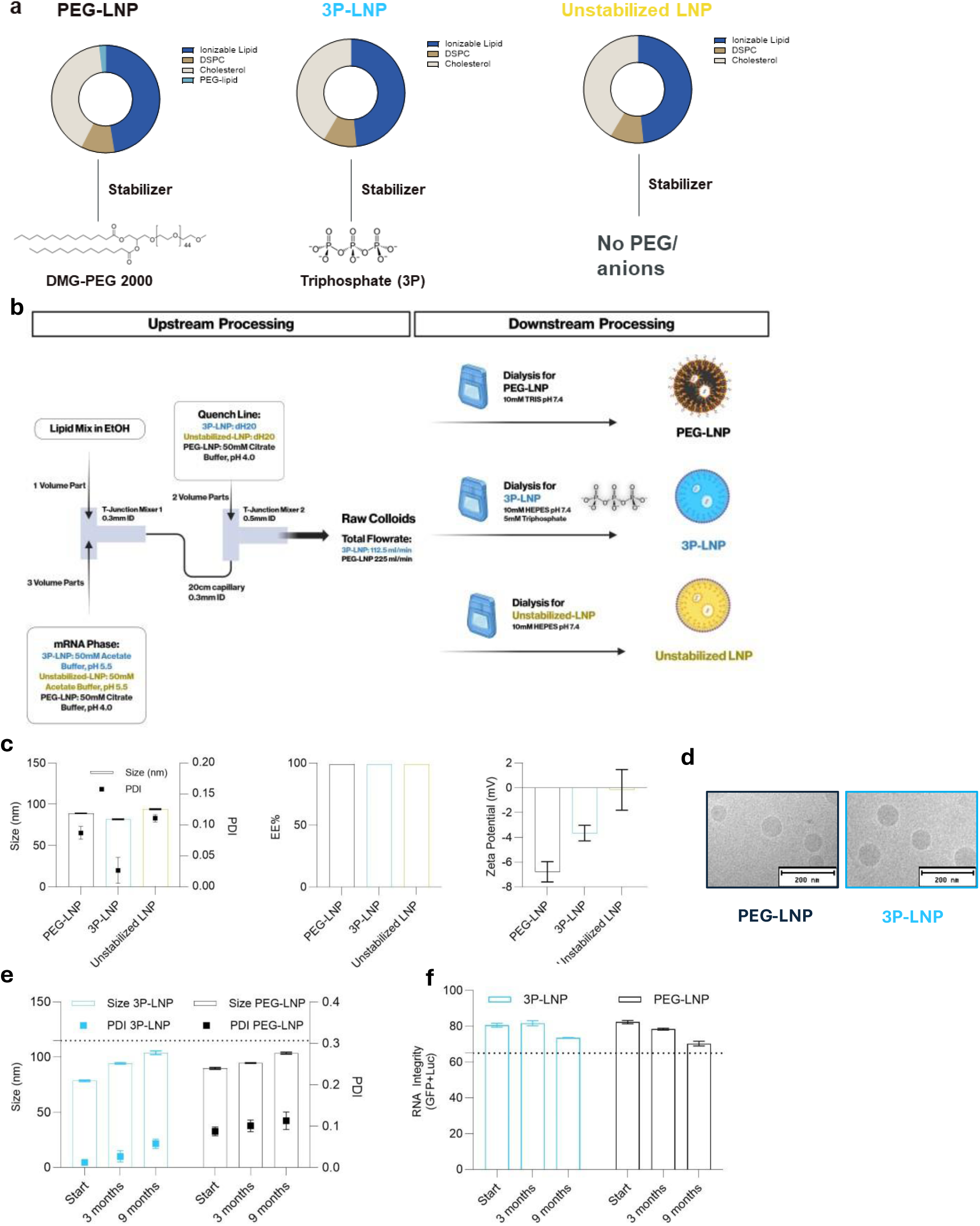
Lipid composition, manufacturing, characterization and stability of PEG-LNPs, 3P-LNPs, and unstabilized LNPs. **(a)** Lipid composition of PEG-LNPs, 3P-LNPs, and LNPs without stabilizers. PEG-LNPs contain DMG-PEG 2000 as a stabilizer, while 3P-LNPs are stabilized with 3P. Unstabilized LNPs lack PEG-lipids or 3P. **(b)** Schematic of the formulation process for PEG-LNPs and PEG-free 3P-LNPs, including upstream and downstream processing steps. Both formulations used a standard ethanol injection protocol. For PEG-LNPs, 50 mM citrate buffer (pH 4.0) was used during the mRNA phase and quenching step. In contrast, 3P-LNPs required 50 mM acetate buffer (pH 5.5) and dH₂O for quenching. Dialysis was performed in 10 mM TRIS buffer (pH 7.4) for PEG-LNPs, whereas 3P-LNPs were dialyzed in 10 mM HEPES buffer (pH 7.4) containing 5 mM 3P. **(c)** Physicochemical characterization of PEG-LNPs, 3P-LNPs, and unstabilized LNPs. (Left) Particle size (Z-average) and PDI of PEG-LNPs, 3P-LNPs, and unstabilized LNPs determined by DLS. All formulations exhibited comparable sizes (∼80 nm) and low PDI (<0.1) after the dialysis step. (Middle) Encapsulation efficiency (EE%) of Luc and GFP mRNA quantified by RiboGreen® assay, showing high efficiency (>99%) for all formulations. (Right) Surface charge (ζ-potential) of PEG-LNPs, 3P-LNPs, and unstabilized LNPs measured by laser Doppler electrophoresis, showing a negative charge between 0 mV and -8 mV. **(d)** Cryo-TEM images of PEG-LNPs and 3P-LNPs reveal spherical morphology, an electron-dense core characteristic of LNPs, and preserved structural integrity. Scale bars, 200 nm. **(e)** Particle size (Size, nm) and PDI of PEG-LNPs and 3P-LNPs measured by DLS. Both formulations remained within the acceptance criteria of ≤115 nm and PDI ≤0.3 throughout the nine-month storage period at 4°C. **(f)** RNA integrity of GFP and Luc mRNA encapsulated in PEG-LNPs and 3P-LNPs measured by fluorescence anisotropy. Both formulations maintained RNA integrity above the defined threshold of 65%, with a slight decline to ∼70% by nine months. Data are presented as mean ± s.e.m. from three technical replicates of the same sample. Abbreviations: 3P, triphosphate; cryo-TEM, cryo-transmission electron microscopy; DLS, dynamic light scattering; EE, encapsulation efficiency; GFP, green fluorescent protein; LNP, lipid nanoparticle; Luc, luciferase; PEG, polyethylene glycol; PDI, polydispersity index; DMG, dimyristoylglycerol; GFP, green fluorescent protein;

The stability study assessed long-term colloidal stability and RNA integrity at 4°C over nine months, with acceptance criteria including the absence of visible aggregates, a particle size ≤115 nm to enable LNP passage through liver sinusoidal endothelial cells (LSECs) [14] and RNA integrity ≥65% to ensure efficient protein translation. Both PEG-LNPs and 3P-LNPs exhibited a modest increase in particle size during storage but remained within the defined criteria for colloidal stability throughout the nine-month period (Fig. 2e). RNA integrity of GFP and Luc mRNA co formulated in 3P-LNPs and PEG-LNPs was monitored over time. Both formulations exhibited high integrity (∼80%) at baseline which was maintained over three months in storage (Fig. 2f). By nine months, a decline in RNA integrity was observed, with both formulations retaining approximately 70% integrity and thereby meeting the criteria for stability. These findings show that replacing PEG with 3P does not affect long-term colloidal stability and RNA integrity. Next, we evaluated the *in vivo* biodistribution, potency, and tolerability of 3P-LNPs compared with PEG-LNPs following a single intravenous administration of 10 µg Luc/GFP mix (1:9 mass ratio) in mice with a group size (n=3) and saline control (n=1). At six hours post-injection, 3P-LNPs elicited approximately two-fold higher signal intensity compared with PEG-LNPs, with the signal of both formulations predominantly localized to the liver (Fig. 3a, b). By 24 hours, signal intensities between the two groups had converged, as confirmed by ex vivo bioluminescence imaging (BLI) (Fig. 3c, d). Mice showed no signs of distress and systemic cytokine and chemokine profiling 24 hours post-injection (Fig. 3e) revealed moderate elevations of interleukin (IL)-6 and tumor necrosis factor alpha (TNF-α) in both 3P-LNP and PEG-LNP groups compared with the saline treated group, consistent with innate immune activation commonly associated with the ionizable lipid [15]. Notably, 3P-LNPs induced higher levels of C-X-C motif chemokine ligand 10 (CXCL10) and monocyte chemotactic protein-1 (MCP-1) compared with PEG-LNPs, suggesting enhanced recruitment of phagocytic cells. To assess hepatotoxicity, serum liver enzyme activity was measured. Mild elevations in alanine aminotransferase (ALT) and aspartate aminotransferase (AST) were observed in the 3P-LNP group, though levels remained within physiologically acceptable ranges when compared to an unrelated cohort of untreated animals (n=19), as indicated by the grey area (Fig. 3f). Lactate dehydrogenase (LDH), a marker of systemic cell damage, showed no significant changes (Fig. 3f). Histopathological analysis of liver tissues using hematoxylin and eosin (H&E) staining revealed intact tissue architecture across all experimental groups, including saline, 3P-LNPs, and PEG-LNPs, with no evidence of necrosis, inflammation, or other significant abnormalities (Supplemental Fig. S4). In summary, 3P-LNPs, lacking PEG-lipids, retained liver-targeted biodistribution without off-target effects and demonstrated a two-fold higher signal intensity at 6 hours post-injection compared with PEG-LNPs, with no observable distress to the mice, significant toxicity or tissue damage. Both 3P-LNP and PEG-LNP formulations triggered mild innate immune activation, with elevated chemokine levels in the 3P-LNP group suggesting phagocytic cell recruitment.

**Figure 3:**
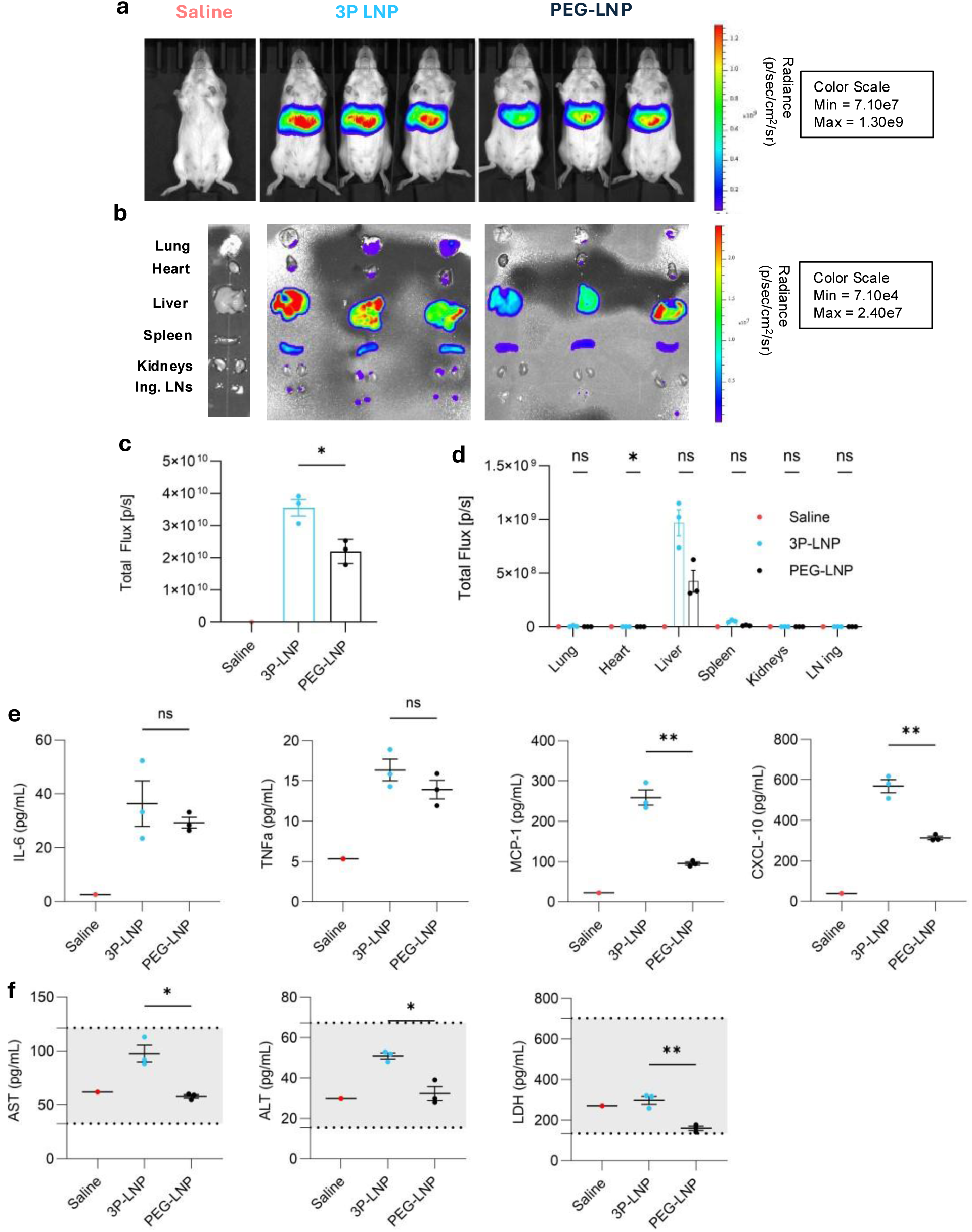
In vivo evaluation of potency, biodistribution and tolerability using bioluminescence imaging (BLI) of 3P-LNPs and PEG-LNPs. **a–d, BLI studies with LNPs. BALB/c mice received a single i.v. injection of saline control (n=1), 3P-LNPs (n=3), or PEG-LNPs (n=3), and Luc expression was monitored at 6 hours post-injection. (a)** Representative whole-body BLI images show strong Luc expression localized predominantly to the liver for both 3P-LNPs and PEG-LNPs, with no signal in the saline group. **(b)** Ex vivo BLI images of major organs confirm predominant hepatic localization of Luc mRNA for both LNP formulations. **(c)** Quantification of total bioluminescent signal in the whole body demonstrates significantly higher Luc activity for 3P-LNPs compared with PEG-LNPs. **(d)** Quantification of bioluminescent signal in specific organs shows enhanced liver-targeted delivery by 3P-LNPs compared with PEG-LNPs. **e–f**, **Evaluation of tolerability and innate immune activation. (e)** Serum cytokine/chemokine levels (TNFα, IL-6, CXCL10, and MCP-1), were measured to assess inflammatory responses. Both 3P-LNPs and PEG-LNPs induced comparable cytokine levels, with 3P-LNPs eliciting higher chemokine levels, indicative of enhanced phagocytic cell recruitment. **(f)** Serum ASAT and ALAT levels were measured to assess hepatotoxicity, showing mild elevations for 3P-LNPs compared with PEG-LNPs, but remaining within physiologically acceptable ranges and no differences were observed for LDH. Levels of naive reference animal cohort (n=19) is indicated by grey shadowed area. For a–f, n = 3 mice per group and n = 1 for the saline in a single experiment. Each symbol represents one animal, and data are presented as mean ± s.e.m. Statistical analysis was performed using an unpaired, parametric two-tailed t-test with Welch’s correction to account for differences in variance between groups. Statistical significance was defined as follows: ns indicates no significance (p > 0.05), * indicates significance (p ≤ 0.05), and ** indicates high significance (p ≤ 0.01). Abbreviations: 3P, triphosphate; ALT, alanine aminotransferase BLI, bioluminescence imaging; CXCL10, C-X-C motif chemokine ligand 10; IL-6, interleukin-6; i.v., intravenous; LNP, lipid nanoparticle; Luc, luciferase; MCP-1, monocyte chemoattractant protein-1; PEG, polyethylene glycol; TNFα, tumor necrosis factor alpha;

The spatial localization and quantitative distribution of GFP⁺ cells in liver tissue were examined by immunofluorescence (IF) staining for GFP, CD68 (Kupffer cell marker), and Endomucin (endothelial marker) (Fig. 4). Both 3P-LNP and PEG-LNP formulations efficiently transfected hepatocytes, producing GFP signals that formed increasing gradients radiating from the central veins toward the lobular periphery. In contrast, the saline control showed no detectable GFP expression. Quantitative analysis revealed similar transfection efficiencies for both formulations, with ∼80 % of liver cells being GFP⁺ (∼2500 cells per mm²) and comparable GFP staining intensities across groups (Fig. 4g). Macrophage abundance in liver tissue varied across groups, with CD68⁺ macrophages comprising 5 % for 3P-LNPs and 15 % for both PEG-LNPs and the saline group (Fig. 4h). The proportion of GFP⁺ CD68⁺ macrophages was higher in the 3P-LNP group compared to PEG-LNPs, suggesting enhanced uptake of 3P-LNPs by liver-resident immune cells.

**Figure 4:**
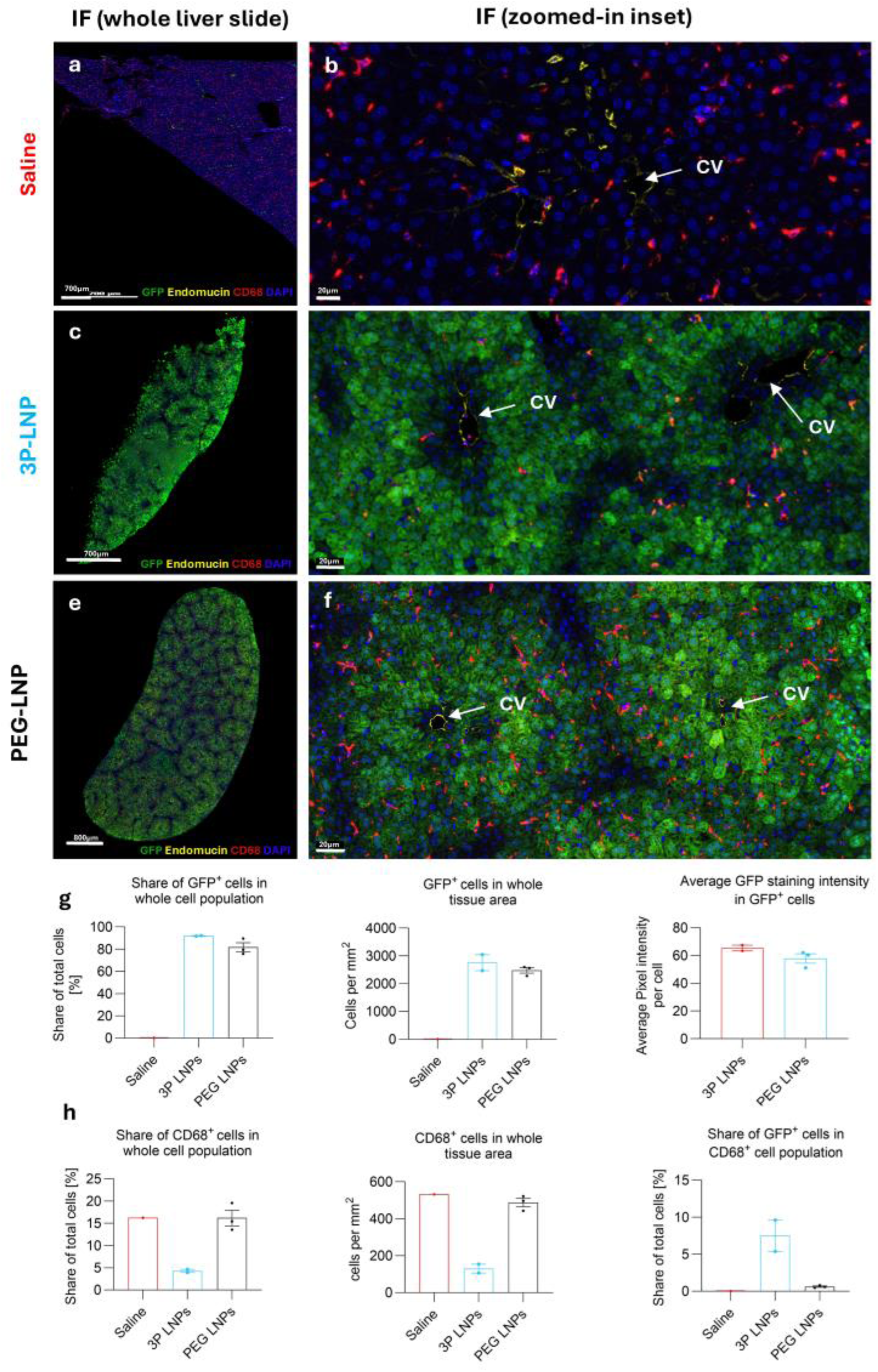
Immunofluorescence analysis of liver tissue following administration of saline, and 3P-LNP, or PEG-LNP formulations, demonstrating GFP expression and localization with endothelium and macrophages. **a-f**, Representative IF images of liver tissue. Panels a, c, and e show whole liver slides, while panels b, d, and f provide zoomed-in insets of the pericentral region surrounding the CV. **(a,b)** Tissue from the saline control (n=1) shows no detectable GFP expression and sparse CD68+ macrophages. 3P-LNP group (n=2) **(c,d)** exhibits strong GFP expression (green) localized to the pericentral region near the CV, with Endomucin+ endothelial cells (yellow) and CD68+ macrophages (red) distributed throughout the tissue. PEG-LNP group (n=3) **(e,f)** shows GFP expression similar to 3P-LNP, localized to the pericentral region, with comparable Endomucin+ and CD68+ cell distributions. The scale bars for image A measure 700 µm, image C; 600 µm, and image E, 800 µm. For the zoomed-in insets within these images, the scale bar is set to 20 µm. **(g)** Quantitative analysis of GFP+ cells. Left: Percentage of GFP+ cells in the total cell population. Middle: Density of GFP+ cells per mm² in the tissue area. Right: Average GFP staining intensity per GFP+ cell. 3P-LNP and PEG-LNP groups show similar GFP expressions. **(h)** Left panel displays the percentage of CD68+ cells within the total cell population. Middle panel shows the density of CD68+ cells per mm² in the tissue area. Right panel illustrates the percentage of GFP+ cells within the CD68+ cell population. Saline-treated and PEG-LNP tissues exhibit similar CD68+ macrophage abundance and lack GFP expression, whereas the 3P-LNP group presents reduced macrophage abundance and increased GFP+ cell presence. Abbreviations: CV, Central Vein; DAPI; 4′,6-diamidino-2-phenylindole; IF, immunofluorescence; GFP, Green Fluorescent Protein;

To distinguish between functional delivery and biodistribution, we employed matrix-assisted laser desorption/ionization mass spectrometry imaging (MALDI-MS Imaging) [16] [17]. Hereby, the spatial localization of key phosphatidylcholine (PC) lipids—PC 32:0, PC 34:1, and PC 38:4 ([M+H]^+^)was analyzed to delineate hepatic microanatomy (Supplementary Fig. S5), all annotated by their fragment ion profiles (Supplementary Figure S6). These endogenous lipids are differentially enriched in central vein (CV), portal triad (PT), and periportal (PP) regions and thus serve as anatomical landmarks [18]. MALDI-MS Imaging provided high-resolution molecular maps of the ionizable lipid BHD-C2C2-PipZ within liver tissue, enabling direct visualization of its spatial distribution after administration of 3P-LNPs or PEG-LNPs (Fig. 5). PC 32:0, localized mainly in CV and PT regions, was co-visualized with BHD-C2C2-PipZ to facilitate orientation. Both formulations exhibited nearly identical total ionizable-lipid levels in the liver, indicating that the absence of PEG did not affect hepatic accumulation—consistent with the comparable GFP expression seen by IF. Notably, MALDI-MS Imaging analysis revealed lower BHD-C2C2-PipZ ion intensities near the CV and higher intensities in PP regions (Fig. 5c), suggesting a spatial distribution that may inversely correlate with GFP expression (Fig. 4).

**Figure 5:**
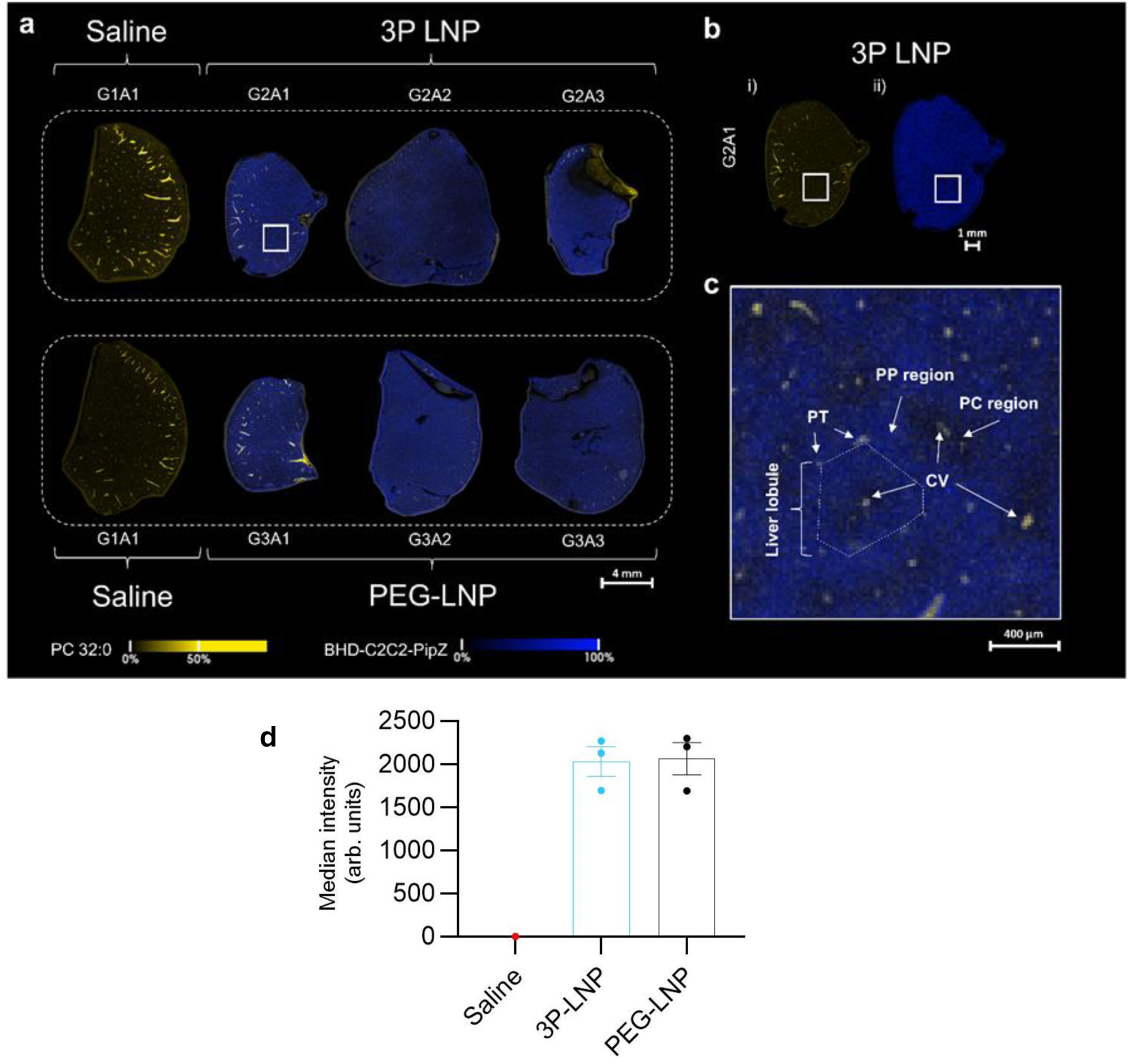
Spatial distribution of the ionizable lipid BHD-C2C2-PipZ in mouse liver via MALDI MS Imaging. **(a)** Merged MALDI MS ion images for PC 32:0 ([M+H]+, m/z 734.570, yellow) and BHD-C2C2-PipZ ([M+H]+, m/z 764.725, blue) for n=3 biological replicates (GNA1 to GNA3, with N=1,2,3) for 3P-LNP and PEG-LNP and saline controls. Ion images were rms normalized and are presented for a mass window of 20 ppm. Macroscopically, the ion intensity for LNPs appears homogeneously distributed across the liver. **(b)** Individual ion images (i, PC 32:0; ii, BHD-C2C2-PipZ) for the tissue section of G2A1. **(c)** Microscopically LNP ion intensity is found more prominently in the PP region and around the (PT compared with the PC region. Arrows indicate the CV and the PC, PP and PT regions. **(d)** Median intensity for BHD-C2C2-PipZ of n=3 biological replicates for the LNP treatment groups. Benjamini-Hochberg corrected p-values were below 0.004 based on a two-sided unpaired t-test compared with the saline treated group. The ratio of the median intensities of both experimental was 0.96(20), consistent with indistinguishable organ specificity for both LNP formulations. Abbreviations: 3P; triphosphate; CV, central vein; LNP, lipid nanoparticles; MALDI MS, matrix-assisted laser desorption/ionization mass spectrometry imaging; PC; pericentral or phosphatidylcholine; PEG, polyethylene glycol; PP, periportal; PT, portal triads; GNA1 to GNA3, with N=1,2,3 representing the different groups and A=1,2,3 the different animals; rms, root mean square.

Our data demonstrates that the inclusion of a stealth lipid, such as a PEG-lipid, may not be as indispensable for LNP performance as previously assumed. Instead, a three-component lipid system can achieve comparable *in vivo* potency, biodistribution, and tolerability without the need for a conventional stealth lipid, challenging a long-standing paradigm in LNP design. By showing that steric shielding can be replaced by electrostatic repulsion, our results establish a new conceptual framework for LNP stabilization. Mechanistically, 3P appears to interact with protonated headgroups of ionizable lipids at the particle surface, forming an anionic layer that provides electrostatic repulsion and may prevent interparticle bridging leading to over nine-month stability. Similar stability was provided by pyrophosphate (2P) consisting of two phosphates (data not shown), whereas monophosphate and citrate were not able to prevent aggregation for unknown reasons (Supplementary Figure S3). Following intravenous administration, 3P-LNPs induced approximately two-fold higher liver transfection at 6 hours post-injection compared to PEGylated controls, while total bioluminescence and GFP expression levels converged after 24 hours (Fig. 3, 4). The enhanced initial transfection likely reflects faster hepatic uptake in the absence of PEG-lipids, which are known to prolong circulation time [4]. This accelerated uptake may transiently increase hepatic nanoparticle flux, resulting in more efficient transgene expression. In contrast, PEG-LNPs might accumulate more gradually, ultimately achieving comparable expression levels after 24 hours. These findings suggest that 3P-mediated stabilization not only preserves biological potency but also accelerates liver transfection. Alternatively, the faster uptake of 3P-LNPs may be attributed to the inhibition of cellular uptake caused by PEG-lipids in LNPs [[19] [20] [21]. Mechanistically, Apolipoprotein E (ApoE) adsorption onto LNPs facilitates hepatic uptake via interactions with low-density lipoprotein receptors (LDL) [22], a process that occurs following the diffusion of PEG-lipids from the particle surface [4]. In contrast, 3P-LNPs, being free of PEG-lipids, do not require this diffusion step for ApoE binding, which may explain the higher signal observed at 6 hours post-injection. The comparable hepatic expression observed with 3P-LNPs indicates that 3P-mediated electrostatic stabilization preserves ApoE binding and receptor-mediated uptake. Mice displayed no signs of distress, and serum levels of IL-6 and TNF-α were comparable between 3P- and PEG-LNP–treated groups. IL-6 and TNF-α are key pro-inflammatory cytokines that mediate innate immune system responses and are often associated with acute inflammation and toxicity [23]. The comparable levels between 3P- and PEG-LNP groups suggest that the absence of a stealth lipid does not influence the safety and tolerability of the LNPs. This challenges the conventional of stealth lipids, which are believed to make nanoparticles invisible to the immune system [24]. Additionally, PEG-lipids were historically used to screen cationic charge and prevent hemolysis [25]; however, as modern LNPs are close to neutral or slightly negative in charge, this function might has become obsolete. These findings suggest that the biological role of PEG-lipids warrants further refinement, highlighting the transformative potential of PEG-free LNPs in advancing lipid nanoparticle technology.

However, chemokines including MCP-1 and CXCL10 were moderately elevated after 3P-LNP administration, consistent with increased transfection of liver-resident CD68+ macrophages. Correspondingly, serum ALT and AST showed small increases relative to PEG-LNP and saline controls, while LDH levels remained unchanged. Together, these findings indicate that mild chemokine and enzyme responses are most likely mediated by the faster hepatic nanoparticle flux of 3P-LNPs rather than intrinsic hepatotoxicity. Immunofluorescence analysis revealed that GFP expression in the liver was predominantly localized around central veins and absent from the periportal region, whereas MALDI imaging showed enrichment of ionizable lipid signals in periportal regions. The finding might be related to hepatic zonation, where periportal hepatocytes primarily support oxidative and lipid metabolism, while centrilobular cells are more active in glycolysis and xenobiotic processing [26]. These data highlight that intrahepatic heterogeneity influences nanoparticle distribution and transgene expression, an important consideration for interpreting biodistribution and potentially optimizing LNP design. Together, our results establish 3P as a functional alternative to PEG for LNP stabilization, combining long-term storage stability with efficient and well-tolerated hepatic delivery. Consistent with previous reports that lowering PEG content to 0.5% enhances uptake by antigen-presenting cells [27], future work should assess whether 3P-LNPs similarly improve antigen presentation and vaccine efficacy and evaluate their performance and pharmacokinetic profiling in repeated dosing regimens.

## Supporting Information

### Formulation of lipid nanoparticles (LNPs)

#### PEG-LNPs

An organic phase comprising an ionizable lipid (di(heptadecan-9-yl) 3,3’-((2-(4-methylpiperazin-1-yl)ethyl)azanediyl)dipropionate), DSPC, cholesterol and DMG-PEG2000 at a molar ratio of 47.5:10:40.7:1.8 was prepared using ethanol as the solvent at a lipid concentration of 47 mM. An RNA phase was prepared with 0.4 mg/mL of RNA in 50mM citrate buffer pH 4.0, diluted from 2 mg/mL RNA in 10 mM HEPES, pH 7.1. As a quenching step, 50mM citrate buffer, pH 4.0, was used. The RNA phase, organic phase, and dilution buffer were continuously mixed in a 3:1:2 (v/v) ratio using a flow rate of 225 ml/min utilizing a Nemesys Flow System (CETONI, Thuringia, Germany) with a mixing T junction with 0.5 mm bore. Formulation volumes ranged from 6 to 20 ml, with the first 6 mL discarded prior to LNP collection. Samples were dialyzed in 2x 12 hours (h) against final buffer (10 mM Tris, pH 7.4) using a Slide A Lyzer dialysis cassette with 10 kDa MWCO dialyzer (Thermo Fisher Scientific, Massachusetts, USA).

#### 3P-LNPs and unstabilized LNPs

An organic phase excluding PEG, comprising lipids of an ionizable lipid (di(heptadecan-9-yl) 3,3’-((2-(4-methylpiperazin-1-yl)ethyl)azanediyl)dipropionate), DSPC and cholesterol at a molar ratio of 48.4:10:41.6 was prepared using ethanol as the solvent at a lipid concentration of 47 mM. An RNA phase was prepared with 0.4 mg/mL of RNA in 50mM acetate buffer, pH 5.5, diluted from 2 mg/mL RNA in 10 mM HEPES, pH 7.4. Distilled water was used for dilution in the quenching step. The RNA phase, organic phase, and dilution buffer were continuously mixed in a 3:1:2 (v/v) ratio using a flow rate of 112.5 ml/min utilizing a Nemesys Flow System (CETONI, Thuringia, Germany) with a mixing T junction with 0.5 mm bore. Formulation volumes range from 6 to 20 ml, with the first 6 mL discarded prior to LNP collection. Samples were dialyzed for 2x 12 h against final buffer of 10mM HEPES, 5mM triphosphate for 3P-LNPs or 10mM HEPES without triphosphate as a negative control (unstabilized LNPs) using a Slide A Lyzer dialysis cassette with 10 kDa MWCO dialyzer (Thermo Fisher Scientific, Massachusetts, USA). For stability experiments and osmolarity adjustments for *in vivo* administration, LNPs samples were added to 1.2M sucrose as cryoprotectant to a final concentration of 300 mM and sterile filtered. Prepared LNPs were stored at 4°C with a final mRNA concentration of 100 µg/ml. Dilutions were performed in the target buffer in the presence of the indicated amount of 3P.

The ionizable lipid was obtained from Croda Pharmaceuticals (Goole, United Kingdom). Cholesterol (no. 700100), DSPC (1,2-distearoyl-sn-glycero-3-phosphocholine, no. 850365), and DMG-PEG 2000 (1,2-dimyristoyl-sn-glycero-3-phosphoethanolamine-N-[methoxy(polyethylene glycol)-2000], no. 880151) were provided by Avanti Polar Lipids (Alabaster, AL, USA).

#### Dynamic Light Scattering (DLS)

The mean particle size and size distribution of LNPs were evaluated using dynamic light scattering (DLS) with a Zetasizer Ultra (Malvern Instruments, Malvern, UK). The instrument utilizes back-scattered light at an angle of 173° to determine particle size. Results are reported as the Z-average particle size (nm) and polydispersity index (PDI). The PDI describes the width of the fitted log-normal distribution around the Z-average size and is calculated using proprietary algorithms within the particle sizing software.

#### Zeta Potential

Zeta potential was measured using laser Doppler velocimetry with a Zetasizer Ultra (Malvern Instruments, Malvern, UK) in folded capillary cells. Values were calculated using the Smoluchowski approximation and expressed in millivolts (mV), providing insights into the surface charge and colloidal stability of the LNPs.

#### Fragment Analyzer (FA) Capillary Gel Electrophoresis

RNA analysis was performed using the Agilent DNF-471 RNA Kit (15 nt) and the 5300 Fragment Analyzer equipped with a 48-capillary array (Agilent Technologies, Santa Clara, CA, USA). The instrument was prepared for analytical runs by loading the required system solutions from the RNA Kit, following the manufacturer’s instructions. LNP samples were first dissolved to release the encapsulated RNA by mixing 20 µL of LNPs (at 0.27 mg/mL or 0.1 mg/mL RNA concentration) with 40 µL of a detergent/solvent mixture containing 20%

Triton X-100 (Sigma-Aldrich, St. Louis, MO, USA) and 30% ethanol (Sigma-Aldrich, St. Louis, MO, USA). The mixture was incubated at 30°C with mild shaking at 600 revolutions per minute (RPM) using an Eppendorf ThermoMixer (Eppendorf, Hamburg, Germany) for 20 minutes. Following dissolution, the samples were diluted 12-fold in Diluent Marker (DM), the Fragment Analyzer loading solution, by adding 12 µL of the dissolved sample to 132 µL of DM in designated wells of a 96-well PCR plate (Thermo Fisher Scientific, Waltham, MA, USA). For the RNA ladder, 2 µL of ladder (Agilent Technologies, Santa Clara, CA, USA) was mixed with 22 µL of DM in a separate well. The plate was sealed with adhesive tape (Thermo Fisher Scientific, Waltham, MA, USA), and the RNA in the samples and ladder was denatured by heating at 70°C for 2 minutes using a heating block (Bio-Rad, Hercules, CA, USA) or PCR cycler (Applied Biosystems, Waltham, MA, USA), followed by immediate cooling on ice. Each denatured sample was divided into three replicates of 48 µL each, which were transferred to adjacent wells for measurement. The prepared plate was loaded into the 5300 Fragment Analyzer (Agilent Technologies, Santa Clara, CA, USA), and samples were injected electrokinetically at 5 kV. Injection parameters were optimized to achieve fluorescence signals within the upper detection range. For samples containing a single RNA species, a final RNA concentration of 7.5 µg/mL (from 0.27 mg/mL LNPs) was combined with an injection time of 5–6 seconds, while a concentration of 2.8 µg/mL (from 0.1 mg/mL LNPs) was injected for 20–25 seconds. RNA separation was performed for 1 hour at 8 kV. Each replicate was evaluated for the presence of a clear peak corresponding to intact RNA, along with low to moderate levels of faster-moving, degraded material. Peak intensity was expected to fall within the range of 5000–60,000 relative fluorescence units (RFU). Peaks were defined by adding drop lines, and the area between the lines was calculated as a fraction of the total signal to determine RNA integrity. A Python-based tool was used for peak analysis to ensure consistent and accurate quantification. This workflow provided reliable and reproducible results for RNA analysis in lipid nanoparticle samples.

### mRNA Synthesis

The luciferase and Green Fluorescent Protein (GFP) mRNA were synthesized using N1-methylpseudouridine (m1Ψ) and CC413, purified using tangential flow filtration, and stored in 10 mM HEPES/0.1 mM EDTA buffer.

### Determination of encapsulation efficiency and encapsulated RNA concentration

Relative encapsulation efficiency and encapsulated RNA concentration were assessed using a Quant-iT RiboGreen assay (Thermo Fisher Scientific, Waltham, Massachusetts). Lipid nanoparticles (LNPs) were diluted to an intra-assay RNA concentration of 0.5 µg/mL in 1× Tris–EDTA (TE) buffer (Thermo Fisher Scientific, Waltham, Massachusetts) and 1× TE buffer supplemented with 1% (v/v) Triton X-100 (Alfa Aesar, Haverhill, Massachusetts). The LNPs were allowed to equilibrate in the buffers for 5 minutes before being transferred to a black 96-well plate (Corning, Corning, New York). A standard curve was prepared following the manufacturer’s instructions, using Luc RNA as the standard. The RiboGreen reagent was mixed 1:1 (v/v) with the standard curve and LNP solution, followed by a 2-minute equilibration. Fluorescence intensity was then measured using a Tecan Spark plate reader (Tecan, Männedorf, Switzerland) at an excitation wavelength of 485 nm and an emission wavelength of 528 nm. RNA content was quantified based on a standard curve generated from a univariate least-squares linear regression, and encapsulation efficiency was subsequently calculated.

### Cryogenic Transmission Electron Microscopy (Cryo-TEM)

In total, 7µl of each sample was applied onto a gold grid covered by a holey Gold Film (UltrAuFoil® 1.2/1.3, Quantifoil^®^ Micro Tools GmbH, Jena, Germany). Excess liquid was blotted automatically from the backside of the grid with a strip of filter paper. Subsequently, the samples were rapidly plunge-frozen in liquid ethane (cooled to –180 °C) in a Cryobox (Carl Zeiss NTS GmbH, Oberkochen, Germany). Excess ethane was removed with filter paper. Samples were transferred immediately with a Gatan 626 cryo-transfer holder (Gatan, Pleasanton, USA) into the pre-cooled Cryo-electron microscope (Philips CM 120, Eindhoven, Netherlands) operated at 120 kV and viewed under low dose conditions. Images were recorded with a 2k CMOS Camera (F216, TVIPS, Gauting, Germany). To minimize noise, four images were recorded and averaged to one image.

### Animal studies

Female 8-week-old BALB/cJRj mice (Janvier Labs, Le Genest-Saint-Isle, France) were selected for the biodistribution and tolerability experimental protocol. Animal husbandry and experimental procedures were performed according to the directives of the EU 2010/63 and were approved by the competent authority of Rhineland-Palatinate, Koblenz, Germany. Mice were intravenously injected into the lateral caudal vein with 100 µL isotonic sodium chlorine solution (0.9% NaCl, Fresenius Kabi, Bad Homburg, Germany) or LNP (test material) containing a total dose of 10 µg mRNA coding for luciferase and eGFP (ratio 1:9). For bioluminescence imaging, 6 hours after intravenous application, the animals were intraperitonially injected with 150 µL D-Luciferin substrate (Revvity Inc., Waltham, MA, USA) at 150 mg/kg bodyweight, followed by imaging under isoflurane anesthesia 5 minutes (min) later using an *in vivo* imaging system (IVIS^TM^ Spectrum, Revvity Inc., Waltham, MA, USA). 24 h after experimental start and 5 min after luciferin administration, final blood samples were drawn from the retro orbital venous plexus. Following euthanasia selected organs including liver and spleen were preserved, imaged and processed for further analysis. Data were analyzed with LivingImage® Software (Revvity Inc., Waltham, MA, USA). Bioluminescence quantification was performed as total flux for the entire mouse at the 6-hour in vivo timepoint and as organ-specific gated flux during ex vivo analysis at 24 hours, expressed as photons per second. Mouse sera were obtained by centrifuging blood samples at 3000 rpm for 20 min at 4 °C. Cytokines and chemokines secretion was measured in mouse sera 24 h after dosing using the V-PLEX custom mouse biomarkers kit (MSD, Maryland, United States), to analyze IL-6, TNF-α, IP-10 and MCP-1 according to the manufacturer’s instructions. The clinical chemistry parameters alanine-aminotransferase (ALT), aspartate-aminotransferase (AST) and lactate dehydrogenase (LDH) were analyzed by Indiko^TM^ Clinical Chemistry Analyzer (ThermoFisher Scientific GmbH, Dreieich, Germany) using its reagent kits according to the manufacturer’s instructions.

### Nuclear magnetic resonance (NMR) Studies

NMR was used to investigate the surface availability of the lipids in the LNP formulation. First the formulation was diluted in the native buffer without cryoprotectant and deuterium oxide (D2O) containing 0.05% trimethylsilylpropionic acid (TMSP) was added to achieve a solution with 10% D2O and formulation concentration of 20 µg/mL in RNA. To characterize the LNP formulation, a 1H-NMR spectra was measured using the pulse programs stebpesgp1s1d (diffusion measurement), which suppresses the signals of water as well as other small molecules via their diffusion properties. For individual lipids, approximately 10 mg of each lipid was dissolved in 0.7 mL deuterated chloroform (CDCl3) containing 0.03% tetramethylsilane (TMS) and pulse program zg30 was performed. All measurements were performed on the NMR spectrometer Avance NEO 600 (Bruker, Karlsruhe, German), magnetic flux density 14.1 Tesla using the TCI cryo probe (Z168380_0016) and automated sample changer Bruker SampleJet.

### Histological analysis

#### Fixation, paraffin embedding, and cutting of organs

Collected organs were fixed for 24 h in 4% ROTI^®^Histofix (Carl Roth, Baden-Württemberg, Germany), cut according to the standardized RITA (Registry of Industrial Toxicology Animal Data) trimming guides and subjected to the standard protocol of dehydration and paraffinization using an ASP300 Tissue Processor (Leica Biosystems, Richmond, Illinois, United States). After embedding in paraffin wax, 3 µm thick sections were prepared using a Leica RM2255 rotatory microtome (Leica Biosystems, Nussloch, Baden-Württemberg, Germany), mounted on SuperFrost^®^ Plus microscopy slides (Fisher Scientific, Hampton, New Hampshire, United States) and baked for 1 h at 56°C in an UN55PA incubator (Memmert, Büchenbach, Bavaria, Germany). Prior to H&E or immunohistochemistry staining, tissue sections were deparaffinized and rehydrated using a Multistainer ST5020 (Leica Biosystems, Nussloch, Baden-Württemberg, Germany) or StainMate Max (Thermo Fisher, Waltham, Massachusetts, United States).

#### H&E staining

Automatic H&E staining was performed with the Multistainer ST5020 (Leica Biosystems, Nussloch, Baden-Württemberg, Germany). Briefly, tissue sections were submitted to nuclear staining with Mayer’s hematoxylin (Carl Roth, Karlsruhe, Baden-Württemberg, Germany) and counterstaining with eosin G (Carl Roth, Karlsruhe, Baden-Württemberg, Germany). Stained sections were dehydrated, and then mounted with CV 5030 Coverslipper (Leica Biosystems, Nussloch, Baden-Württemberg, Germany).

#### Triple IF staining

Baked slides were deparaffinized and rehydrated in Multistainer ST5020 (Leica Biosystems, Nussloch, Baden-Württemberg, Germany). Next, heat-induced epitope retrieval was performed in 1X AR9 buffer (Akoya Biosciences, Marlborough, Massachusetts, United States; # AR900250ML) using a CertoClav Multicontrol 12L Alu sterilizer (CertoClav Sterilizer GmbH, Traun, Austria) for 10 min at 120°C, followed by a 10 min cooling period at room temperature (RT). Slides were transferred to the Discovery Ultra Multistainer (Roche Diagnostics, Penzberg, Bavaria, Germany), where they were quenched with endogenous peroxidase for 16 min at RT using Discovery Inhibitor (Roche Diagnostics, Penzberg, Bavaria, Germany; # 07017944001), followed by application of a rabbit anti-GFP antibody (Abcam, Cambridge, Cambridgeshire, United Kingdom; clone EPR14104, #ab183734, 1:100 dilution in 10% goat serum) at 37°C for 60 min. OmniMap HRP-anti-rabbit secondary antibody (Roche Diagnostics, Penzberg, Bavaria, Germany; #5269679001) was applied at 37°C for 32 min. Primary–secondary antibody complex was visualized using Opal520 (Akoya Bioscience, Marlborough, Massachusetts, United States; FP1487001KT) as a substrate for the horseradish peroxidase (HRP), diluted 1:250 in 1x Plus Amplification Reagent (Akoya Bioscience, Marlborough, Massachusetts, United States; # FP1609). For a second round of staining, a stripping step with Reaction Buffer (Roche Diagnostics, Penzberg, Bavaria, #05353955001) at 95°C for 12 min was applied, followed by rabbit anti-endomucin (Abcam, Cambridge, Cambridgeshire, United Kingdom; clone V.7C7.1, #ab106100; 1:250 dilution in 10% goat serum) and secondary antibody staining with OmniMap HRP-anti-rabbit antibody. Staining was visualized using Opal^TM^570 (Akoya Bioscience, Marlborough, Massachusetts, United States; #FP1488001KT; 1:400 dilution in 1x Plus Amplification Reagent). Final staining involved a stripping step at 95°C for 12min followed by incubation with Reaction Buffer with rabbit anti-CD68 (Cell Signaling Technologies, Danvers, Massachusetts, United States; clone E3O7V, #97778; dilution of 1:2000 in 10% goat serum) for 60 min at 37°C and Omnimap anti-rabbit HRP as secondary antibody at 37°C for 32 min. Staining was visualized using Opal^TM^690 (Akoya Bioscience, Marlborough, Massachusetts, United States; # FP1497001KT, 1:400 dilution in 1x Plus Amplification Reagent) for 20 min at RT. Lastly, tissues were counterstained with (4’,6-diamidino-2-phenylindole) (DAPI) (Sigma Aldrich, St. Louis, Missouri, United States; #102352760; 1:2,500 in phosphate buffered saline), before slides were washed with TBST and mounted using fluorescence mounting medium (DAKO, Glostrup, Capital Region, Denmark; #S3023).

#### Image acquisition

Following H&E, IHC, and triple IF stainings, the dried and mounted sample slides were scanned with the NanoZoomer S360 (Hamamatsu Photonics, Shizuoka, Japan) with a 20× objective (H&E, IHC), or with the PhenoImager HT (Akoya Biosciences, Marlborough, Massachusetts, United States; triple IF) with a 20× objective, and their respective device software.

#### Cell Counting / Digital Image Analysis

Digital image analysis was conducted using HALO software (version 4.1; Indica Labs, Albuquerque, NM, USA). Liver tissue regions were automatically identified through deep learning-based tissue segmentation algorithms integrated within the HALO platform. Artifacts, such as dirt and tissue folds, were excluded using a one-shot segmentation tool, ensuring accurate tissue delineation. CD68-positive areas were segmented using a pre-trained dense residual neural network, optimized for immunohistochemical marker detection. GFP-positive regions were quantified using the Indica Labs Area Quantification FL (version v3.0.1). Liver parenchymal cells were segmented and GFP mean fluorescence intensity measured using the Indica Labs HighPlex FL algorithm (version 5.2.2).

#### Matrix Assisted Laser Desorption/Ionization (MALDI) mass spectrometry imaging

Liver sections of 10 µm thickness were collected at –15° C using a CM1860 UV Cryostat (Leica Biosystems, Nussloch, Germany) and mounted onto indium tin oxide (ITO)-coated glass slides (Bruker Daltonics, Bremen, Germany).

MALDI matrix was applied with an M5 sprayer (HTX Technologies, Chapel Hill, NC, USA) using 10 mg/mL 2,5-dihydroxyacetophenone (DHAP) dissolved in Acetonitrile:water (70:30). The spray parameters were set as follows: 10 psi nitrogen pressure, nozzle temperature of 75 °C, tray temperature of 30 °C, 10 passes with a flow rate of 0.1 mL/min at 1200 mm/min velocity and 3 mm track spacing.

MALDI MS imaging was conducted on a timsTOF fleX mass spectrometer (Bruker Daltonics, Bremen, Germany) in positive-ion mode with a 200 shots per pixel using a laser spot size and a raster width of 20 µm each. Transfer parameters were set as follows: MALDI Plate Offset= 50.0 V, Deflection 1 Delta= 70.0 V, Funnel 1 radio frequency (RF) = 350.0 Voltage Peak-to-Peak (Vpp), in-source Collision-Induced Dissociation (isCID) Energy= 0.0 eV, Funnel 2 RF= 350 Vpp, Multipole RF= 300 Vpp. Collision cell energy was set to 10.0 eV, and the collision RF was set to 1800.0 Vpp. The quadrupole was set to an Ion Energy of 5.0 eV and a Low Mass of 300 m/z. Focus Pre TOF Transfer time was set to 85.0 µs, the Pre Pulse Storage was set to 10.0 µs.

*m/z* features of interest were MS2-fragmented by imaging parallel reaction monitoring-parallel accumulation serial fragmentation (iprm-PASEF) [28] using the following trapped ion mobility spectrometry (TIMS) settings: 1/K_0_ Start= 0.70 V*s/cm2, 1/K_0_ End= 1.80 V*s/cm2, Ramp time= 200 ms, Accu. Time= 20.1. Transfer parameters were set as follows: MALDI Plate Offset= 50.0 V, Deflection 1 Delta= 70.0 V, Funnel 1 RF= 350.0 Vpp, isCID Energy= 0.0 eV, Funnel 2 RF= 350 Vpp, Multipole RF= 300 Vpp. Collision cell energy was set to 25.0 eV, and the collision RF was set to 800.0 Vpp. The quadrupole was set to an Ion Energy of 5.0 eV and a Low Mass of 300 m/z. Focus Pre TOF Transfer time was set to 75.0 µs, the Pre Pulse Storage was set to 10.0 µs. The isolation window was set to m/z 1.5. The collision energy gradient from *m/z* 500 to 900 to 1200 was set to 30, 50, and 65 eV, respectively.

Data were acquired using tims Control V6.1.1 and flexImaging V7.6. Data analysis was performed using SCILS Lab V 2025b Pro (all Bruker Daltonics). To directly compare relative abundancies of the LNPs in liver, the data were exported from SCILS Lab as imzmL data format and analyzed in python using pyM^2^aia [29] and common statistical analysis

## Acknowledgements

We also thank Aykut Zelak for providing the cyto- and chemokine data. Rieke Meffert and Diana Schneider for support in animal studies and providing clincial chemistry data. Nadine Salisch for reviewing the manuscript with focus on immunology. Many thanks also to the analytical team: Daniel Cassier, Damla Cetin, Jorrit-Jan Krijger and Ankush Sood. The schematic illustrations were created using BioRender.com.

## Author contributions: CRediT

**Gholan amraz:** Conceptualization, Methodology, Writing - Original draft preparation, Formal analysis, Investigation, Visualization, Project administration

**Mario Saucedo-Espinosa:** Conceptualization, Supervision, Writing - Review & Editing

**Kerstin Walzer:** Resources, Investigation, Formal analysis, Visualization

**Florian Frohns:** Investigation, Formal analysis, Visualization

**Elfi Katrin Schlohsarczyk:** Investigation, Formal analysis

**Jakob Käppler:** Formal analysis, Visualization

**Björn Fröhlich:** Investigation, Formal analysis, Visualization

**Stefan Schmidt:** Investigation, Formal analysis, Visualization, Writing - Review & Editing

**Carsten opf:** Formal analysis, Supervision, Writing - Review & Editing

**Steffen Panzner:** Conceptualization, Methodology, Supervision, Writing - Review & Editing

## Conflicts of Interest Disclosure

| Author name | COI |
| --- | --- |
| Gholan Hamraz | Gholan Hamraz is an employee of BioNTech SE and may hold stock or stock options |
| Mario Saucedo-Espinosa: | Mario Saucedo-Espinosa is an employee of BioNTech SE and may hold stock or stock options |
| Kerstin Walzer | Kerstin Walzer is an employee of BioNTech SE and may hold stock or stock options |
| Florian Frohns | Florian Frohns is an employee of BioNTech SE and may hold stock or stock options |
| Elfi Katrin Schlohsarczyk | Elfi Katrin Schlohsarczyk is an employee of BioNTech SE and may hold stock or stock options |
| Jakob Käppler | Jakob Käppler is an employee of BioNTech SE and may hold stock or stock options |
| Björn Fröhlich | Björn has no conflicts of interest to declare. |
| Stefan Schmidt | has no conflicts of interest to declare. |
| Carsten Hopf | Carsten Hopf has no conflicts of interest to declare. |
| Steffen Panzner | Steffen Panzner is an employee of BioNTech SE and may hold stock or stock options |

## Supplementary Material

**Figure S1:**
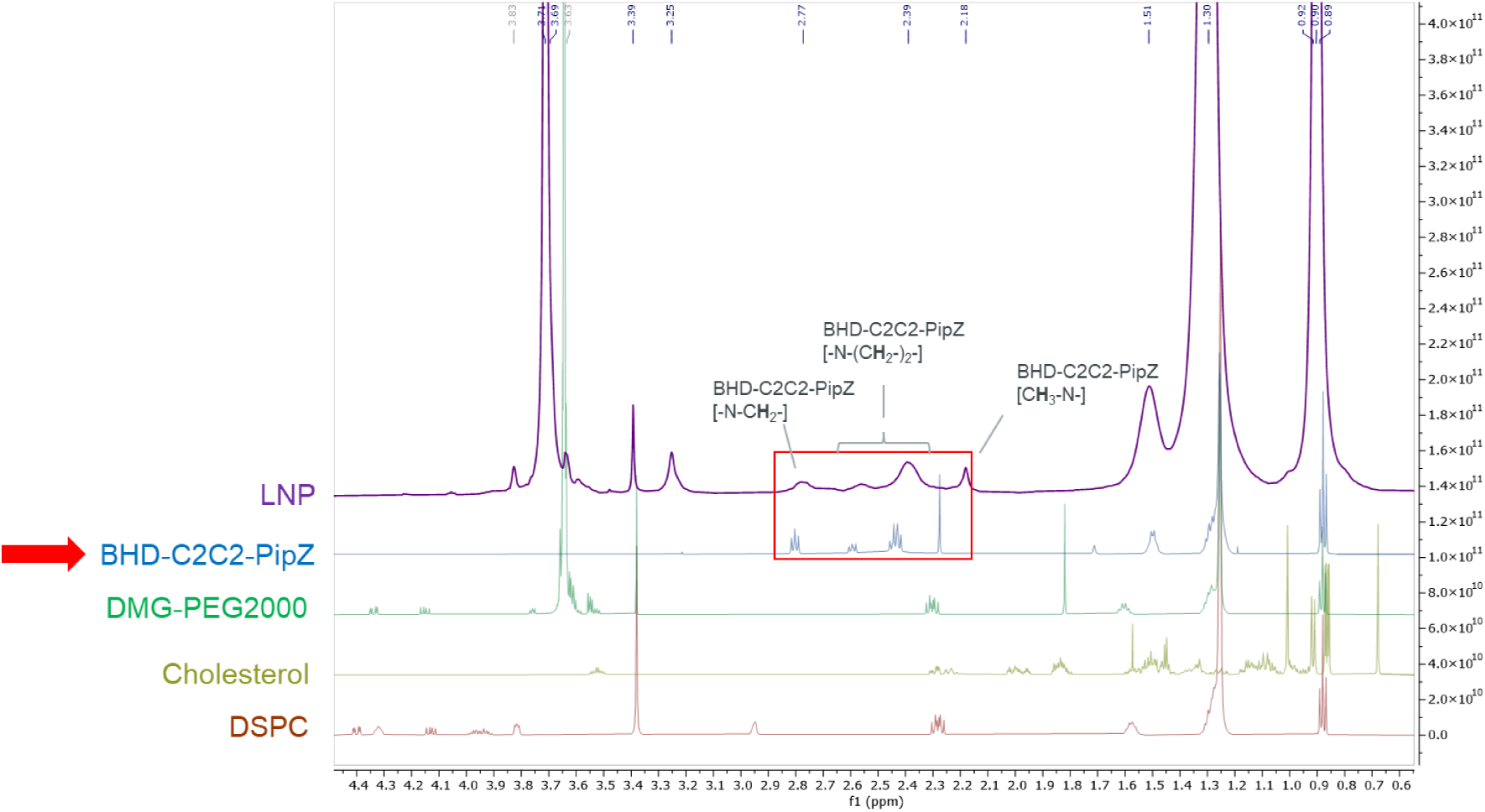
Surface Proton nuclear magnetic resonance (1H-NMR) spectra comparing the lipid nanoparticle (LNP) formulation (purple) with its individual components. Surface proton nuclear magnetic resonance (1H-NMR) spectra comparing the lipid nanoparticle (LNP) formulation (purple) with its individual components: BHD-C2C2-PipZ (blue), DMG-PEG2000 (green), cholesterol (yellow), and DSPC (red). NMR experiments were performed using the pulse program stebpesgp1s1d for the LNP formulation and zg30 for individual lipids on a Bruker Avance NEO 600 spectrometer with a TCI cryo probe. Key spectral features corresponding to specific chemical groups are annotated, highlighting the molecular contributions of each component to the LNP surface structure.

**Figure S2:**
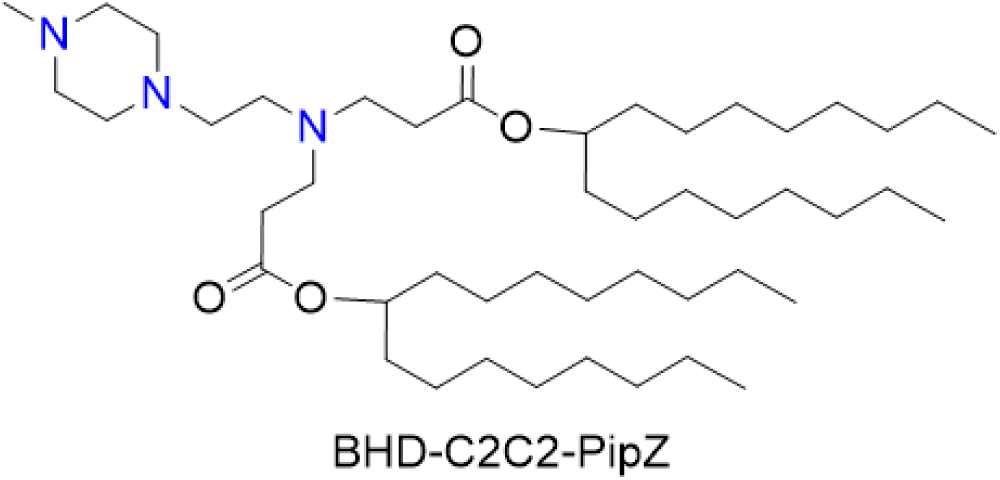
Chemical structure of the ionizable lipid BHD-C2C2-PipZ, designed for lipid nanoparticle (LNP) formulations.

**Figure S3:**
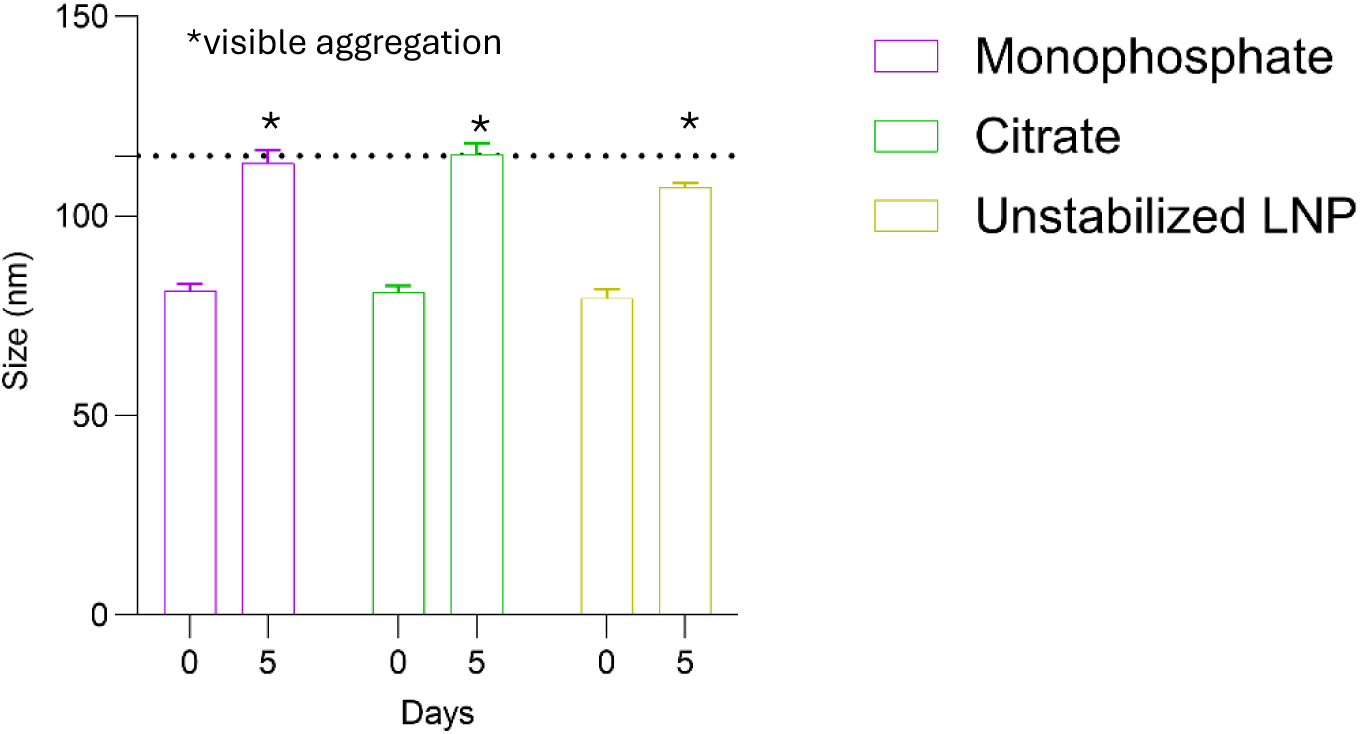
Colloidal stability of unstabilized LNPs and formulations containing monophosphate or citrate over 5 days. Particle size (nm) of lipid nanoparticles (LNPs) measured at Day 0 and Day 5 using dynamic light scattering (DLS). Formulations containing monophosphate (purple), citrate (green), or **u**nstabilized LNPs (yellow) exhibited significant increases in particle size over 5 days, exceeding the acceptance criterion of ≤115 nm (dotted line). Asterisks (*) indicate formulations with visible aggregation at Day 5. Data are presented as mean ± s.e.m. from three technical replicates of the same sample (n = 3). Abbreviations: LNP, lipid nanoparticle; DLS, dynamic light scattering.

**Figure S4:**
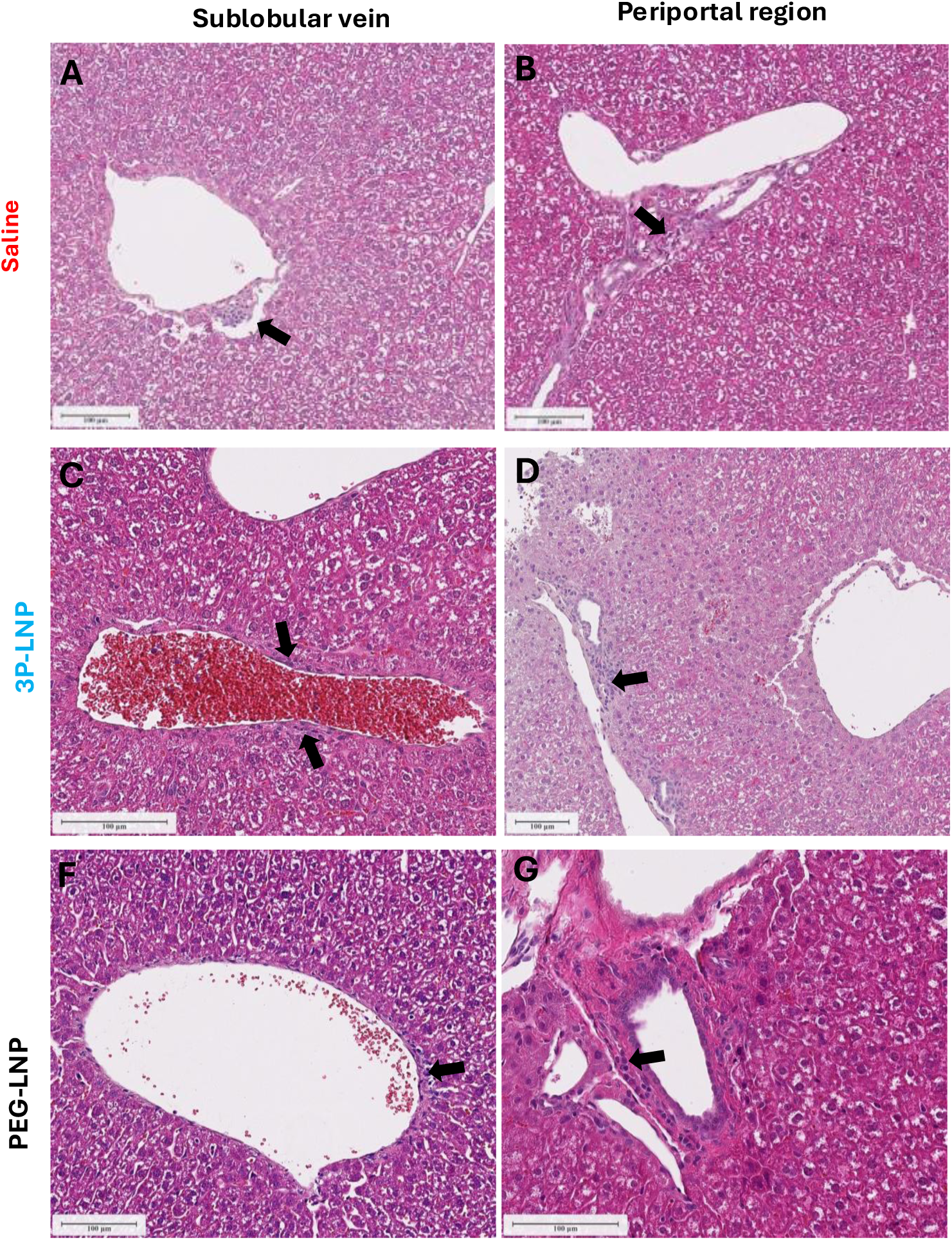
Representative histopathological findings in livers from mice after intravenous administration of saline or LNP formulations (H&E staining). Livers from animals treated with (A, B) Saline, (C,D) 3P-LNPs (10 µg, Luc/eGFP 1:9), and (E,F) PEG-LNPs (10 µg, Luc/eGFP 1:9) showed minimal histopathological changes. Single mixed inflammatory infiltrates composed of mononuclear cells and neutrophilic granulocytes (arrows) were occasionally observed around sublobular veins and periportal regions across all groups. Taken together, these findings were considered as background changes and not related to the test items. Scale bars: 100 µm

**Figure S5:**
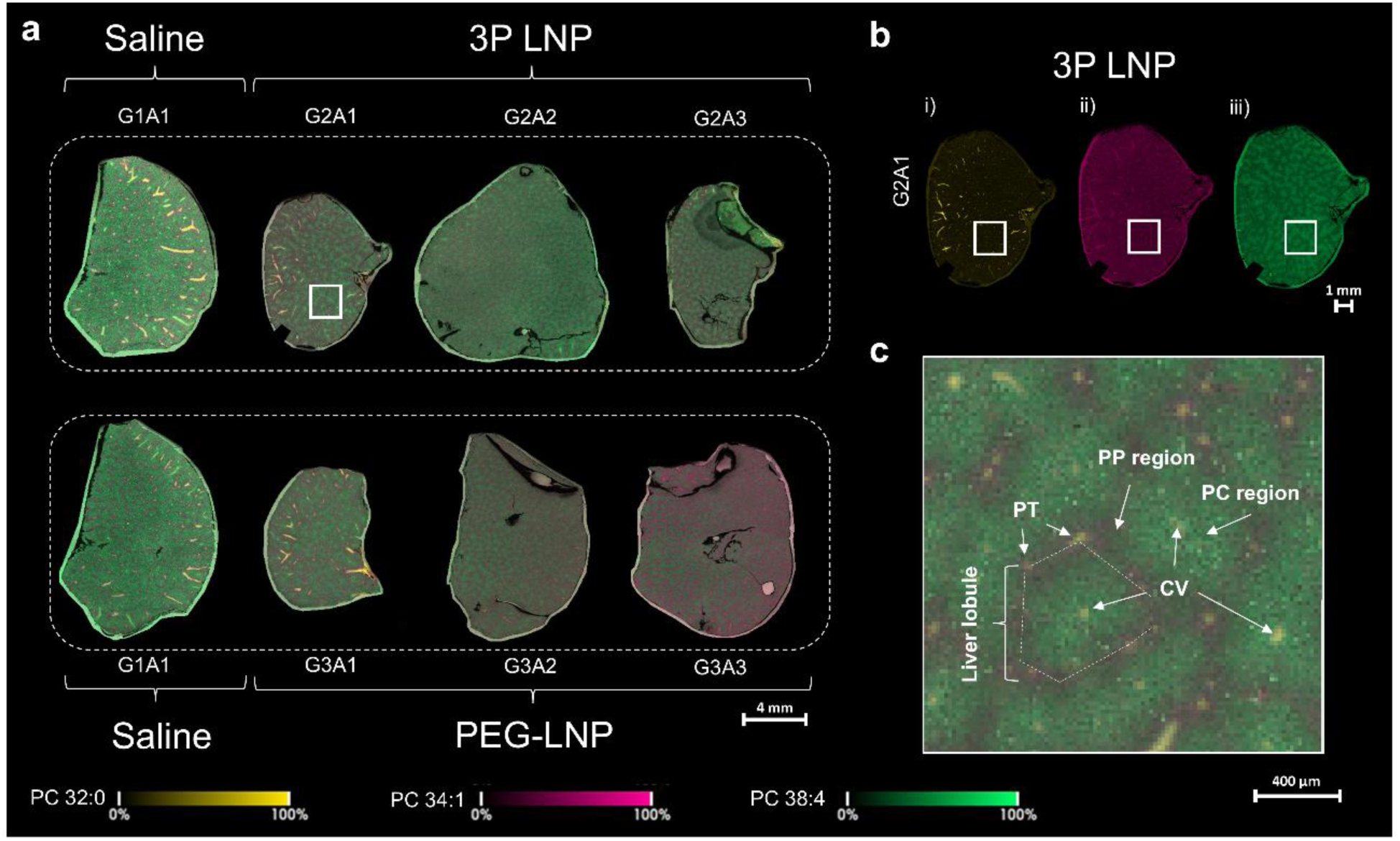
Lipid heterogeneity mimic the hepatic parenchyma in mouse liver via MALDI MSI. **(a)** Composed ion images for PC 32:0 ([M+H]+, m/z 734.570, yellow), PC 34:1 ([M+H]+, m/z 760.587, magenta) and PC 38:4 ([M+H]+, m/z 810.602, green) for the treatment groups G1 (saline), G2 (3P LNP) and G3 (PEG-LNP). Data are presented for n=3 biological replicates (GNA1 to GNA3 with N=1,2,3) for G2 and G3. Ion images are presented for a mass window of 20ppm. MALDI MSI data was rms (root mean square) normalized and was acquired on a timsTOF flex instrument. **(b)** Individual ion images (i, PC 32:0; ii, PC 34.1; iii, PC38.4) for the tissue section of G2A1. In agreement with [18]. PC 32:0, PC34:1, and 38:4 are primarily abundant in PTs/CVs and PP region, respectively. **(c)** Functional structures of the liver lobule are indicated in the overlayed ion image, highlighting the periportal (PP) region surrounding the portal triad (PT) and the pericentral (PC) region surrounding the central vein (CV).

**Figure S6:**
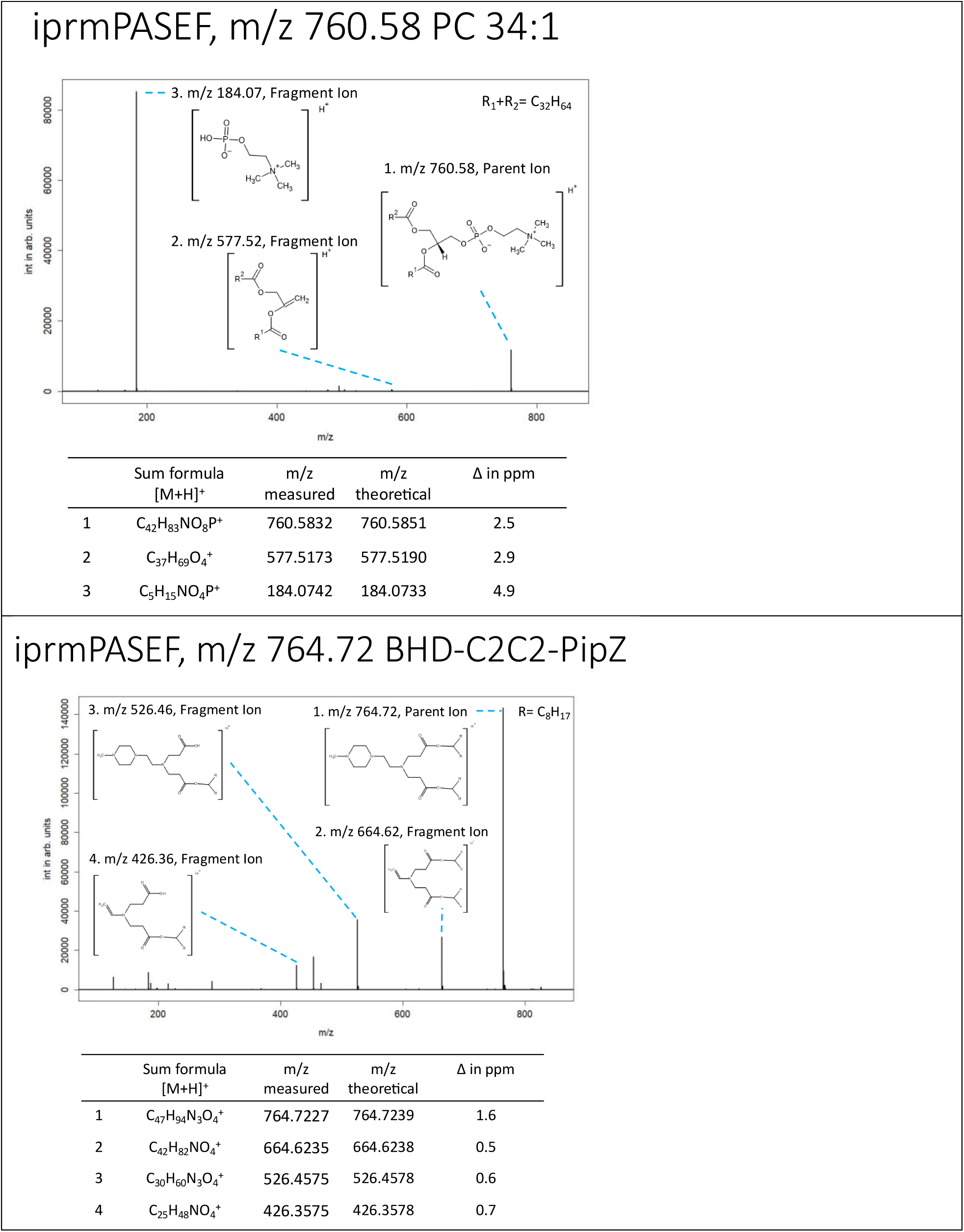
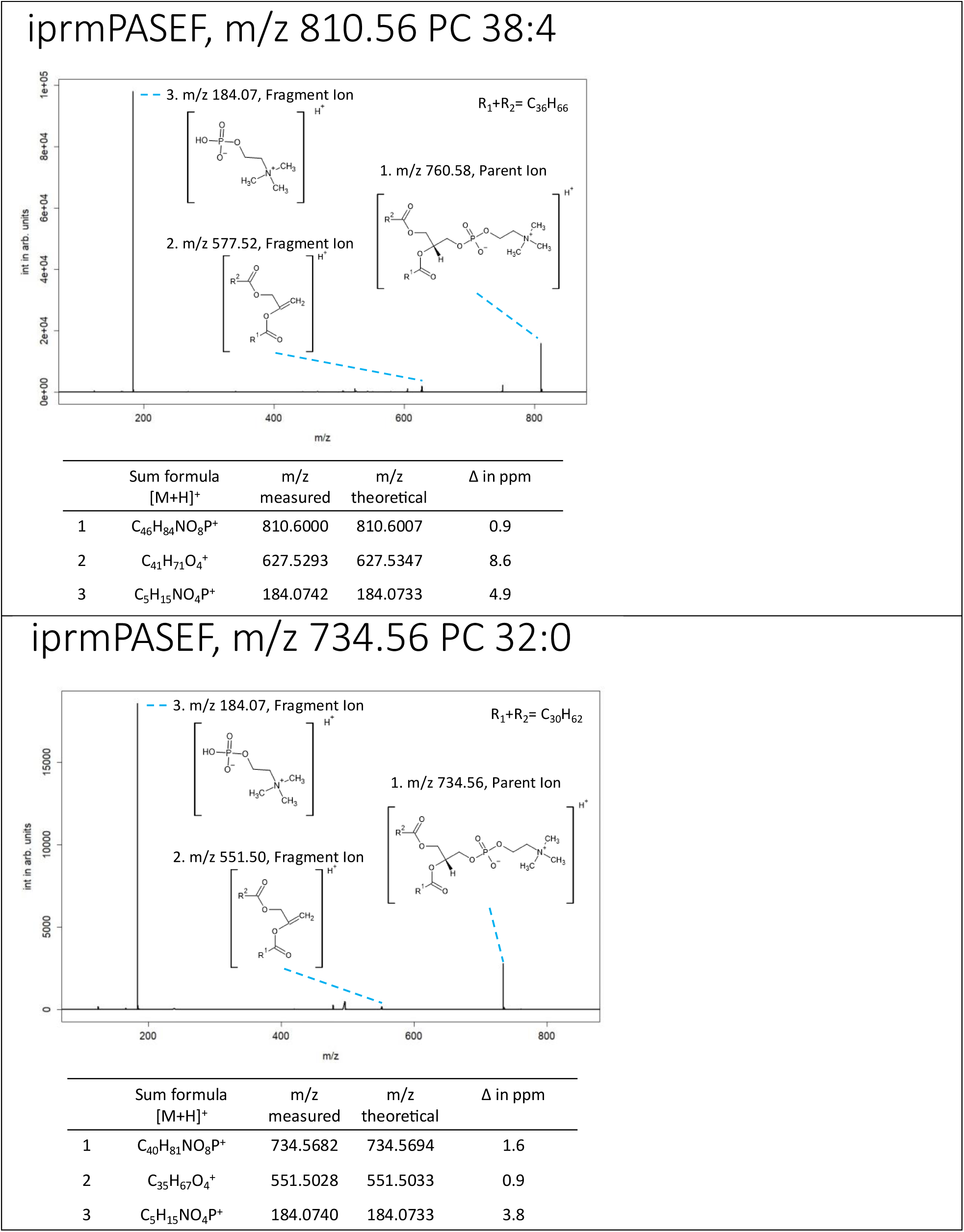
Fragment Ion Analysis of Lipids Used for MALDI-MSI. The table summarizes the fragment ion profiles of key lipids detected in MALDI-MSI, including PC 34:1, PC 38:4, PC 32:0, and the ionizable lipid BHD-C2C2-PipZ. Each lipid is characterized by its parent ion (m/z) and corresponding fragment ions, with measured and theoretical m/z values, molecular formulas, and mass accuracy (Δm/z in ppm). Fragmentation patterns provide structural insights into lipid composition and localization within hepatic tissue. These lipids were selected to reflect hepatic parenchymal structure, with PC species primarily abundant in regions around the central vein (CV), portal triads (PTs), and periportal (PP) areas. The ionizable lipid BHD-C2C2-PipZ was co-imaged with PC 32:0 to describe spatial distribution in detail. in ppm). Fragmentation patterns provide structural insights into lipid composition and localization within hepatic tissue. These lipids were selected to reflect hepatic parenchymal structure, with PC species primarily abundant in regions around the central vein (CV), portal triads (PTs), and periportal (PP) areas. The ionizable lipid BHD-C2C2-PipZ was co-imaged with PC 32:0 to describe spatial distribution in detail.

